# Meta-analyses of *Culex* blood-meals indicates strong regional effect on feeding patterns

**DOI:** 10.1101/2024.05.24.595669

**Authors:** Jet S. Griep, Eve Grant, Jack Pilgrim, Olena Riabinina, Matthew Baylis, Maya Wardeh, Marcus Blagrove

**Author notes:** = corresponding authors.

## Abstract

Understanding host utilization by mosquito vectors is essential to assess the risk of vector-borne diseases. Many studies have investigated the feeding patterns of *Culex* mosquitoes by molecular analysis of blood-meals from field collected mosquitoes. However, these individual small-scale studies only provide a limited understanding of the complex host-vector interactions when considered in isolation. Here, we analyse the *Culex* blood-feeding data from 90 publications over the last 15 years to give a global insight into the feeding patterns of *Culex* mosquitoes, with particular reference to vectors of currently emerging *Culex*-borne viruses such as West Nile and Usutu. Data on 26,857 blood-meals from 71 different *Culex* species were extracted from published literature. The percentage of blood-meals on amphibian, avian, human, non-human mammalian, and reptilian hosts was determined for each *Culex* species. Our analysis showed that feeding patterns were not significantly explained by mosquito species-level phylogeny, indicating that external factors play an important role in determining mosquito feeding patterns. For *Cx. quinquefasciatus, Cx. pipiens* complex, and *Cx. tritaeniorhynchus*, feeding patterns were compared across the world’s seven biogeographical realms. *Culex tritaeniorhynchus, Cx. pipiens* complex and *Cx. quinquefasciatus* all had significantly varied feeding patterns between realms. These results demonstrate that feeding patterns of *Culex* mosquitoes vary between species but can also vary between geographically distinct populations of the same species, indicating that regional or population-level adaptations are major drivers of host utilization. Ultimately, these findings support the surveillance of vector-borne diseases by specifying which host groups are most likely to be at risk.

**Author summary:** Being aware of mosquito biting behaviour is essential to determine the threat of mosquito-borne diseases. Studying the feeding patterns of *Culex* mosquitoes is crucial as these mosquitoes are vectors of currently emerging or re-emerging arboviruses such as West Nile and Usutu. Feeding behaviour of *Culex* mosquitoes has been examined in many individual small-scale studies. These studies only focus on the feeding patterns in a specific area. To gain a more global understanding of these feeding patterns we analysed all available *Culex* blood-feeding data from the last 15 years. In summary, data on 26,857 blood-meals from 71 different *Culex* species was collected. For each species the percentage of blood-meals on different host groups was determined. We analysed the relationship between feeding patterns and mosquito phylogeny, which showed that phylogeny alone could not explain feeding patterns. These results indicate that external factors such as land use and climate could play an important role in determining feeding patterns. For further analysis we determined the feeding patterns for three important vector species, *Cx. quinquefasciatus, Cx. pipiens* complex, and *Cx. tritaeniorhynchus* in different biogeographical realms. All three species showed different feeding patterns in the included realms. Thus, the same species can have different feeding patterns in different regions, indicating the importance of local surveillance.

## Introduction

Most mosquito species are restricted to either tropical or temperate areas, but mosquito species of the *Culex* genus occur in both (1,2). *Culex* mosquitoes can be vectors for viruses such as West Nile virus, Japanese encephalitis virus, Rift valley fever virus, and several others (3–5). Many *Culex* species can act as reservoir vectors, transmitting pathogens between reservoir hosts, or bridge vectors, transmitting from reservoir hosts to dead-end hosts which, in some cases, includes humans. Several *Culex* species have been implicated in the transmission of emerging and re-emerging diseases such as West Nile and Usutu (6–8).

Assessing the feeding patterns of mosquito vectors is essential in determining the epidemic potential of certain mosquito-borne pathogens. The hosts that mosquitoes have fed on can be determined by molecular analysis of the blood-meals in the midguts of field collected mosquitoes (9). If there is a high percentage of both human and reservoir blood-meals and an overlap in time and space of the vector and the host, the risk of efficient pathogen transmission to humans increases significantly (10). Additionally, as feeding patterns can shift due to both climate and land use change, it is important to be aware of recent feeding patterns (11,12).

Both intrinsic and extrinsic factors can influence feeding patterns. Intrinsic factors such as genetics can affect host preference (13), but do not fully explain feeding patterns. For example, despite having limited genetic differences *Culex pipiens* form *pipiens* and *Culex pipiens* form *molestus* have very distinct feeding patterns. Form *pipiens* is believed to feed more on birds whilst form *molestus* is believed to mostly utilize mammalian hosts. The mechanism underlying this difference in feeding patterns is unknown (14), but is likely influenced by external factors. Examples of these are host availability and seasonality (15). Host defensive behaviours and learning could also influence host utilization. If a mosquito for example had a successful feed on a host it is more likely to go for the same host species for the second feed (16). Since both intrinsic and extrinsic factors can play a role in host selection of mosquitoes the exact mechanism behind this is hard to elucidate.

Many species of the *Culex* genus are able to feed on a wide variety of hosts (17). *Culex* feeding patterns have been studied frequently in small-scale individual studies, focussing on the regional feeding patterns of the mosquitoes from that area. As feeding patterns are affected by a variety of external factors it is expected that they might also differ between biogeographic realms. These realms represent an area that has a similar plant an animal distribution and that experienced a distinct evolutionary history and climate (18). As the realms have different assemblages of animals and plants; but share certain mosquitoes, we wanted to explore feeding patterns of important vector species in these realms. This could show if different realms require specialized surveillance strategies.

Here, we analyse the data published over the last 15 years describing *Culex* blood-feeding in the field. This analysis enables a global insight into the feeding patterns of *Culex* mosquitoes and thus provides higher-level data for risk assessments to specific hosts, and to specific regions, which will inform ongoing efforts, particularly in Europe (19,20), to mitigate the spread of *Culex*-borne viruses (e.g. WNV and USUV); as well as assessing the risk of the spread of invasive *Culex* vectors e.g. *Cx. modestus* (21).

## Methods

### Data collection

#### Meta-analysis

*Culex* blood-feeding studies were collected using the PubMed database. For the initial search the keywords: “*Culex*” and “Feeding-patterns” were utilized to collect review papers, systematic reviews, meta-analyses, and primary papers containing blood-meal analyses of *Culex* mosquitoes. Following this initial review, the combination of keywords: “*Culex*”, “blood meal”, “bloodmeal”, “blood-meal”, “mosquito”, “host-species”, “host”, and “feeding” was used. For several species, where data were lacking, additional searches were performed included the full species name. English literature from the time period from 2008-2023 was included in this review. This 15-year time period was chosen to focus on recent feeding patterns of *Culex* mosquitoes. Papers were included in our analysis if they utilized molecular methods to confirm the blood-meal source. Studies which analysed the presence of *Culex* species in animal traps and laboratory-based blood-feeding studies were excluded. Studies performed in zoos were included.

From the articles that were included the following information was extracted: *Culex* species, mosquito collection location, realm, country, vertebrate host species, location of vertebrate host species, larger host group classification (amphibian, avian, human, non-human mammalian, and reptilian), fed (yes/no), number of blood-meals on host, total number of blood-meals recorded for a specific species in one study, the percentage of blood-meals from one mosquito on one host compared to all recorded blood-meals for that mosquito species in a specific study. The realm was determined based on the Udvardy (1975) system. If the specific form of *Culex pipiens* was not specified, the data was categorized under *Culex pipiens sp*. Furthermore, for our analysis *Culex pipiens sp*., *Culex pipiens molestus, Culex pipiens pallens, Culex pipiens pipiens, Culex pipiens/molestus* hybrid, *Culex pipiens/torrentium* were clustered as “*Culex pipiens* pooled”. In the dataset they were all kept separate if they were recorded to the biotype level in the original paper. The hosts were described at the level recorded in literature. All data collected for this study and the code used to analyse the data are available in the supporting information (S1 and S2).

### Data analysis

#### Feeding patterns

The number of blood-meals per host species was recorded. All hosts were aggregated into five main groups: amphibian, avian, human, non-human mammalian, and reptilian. Based on the total number of blood-meals recorded for each species the percentage of blood-meals from each of these five groups was calculated. All calculations and data visualizations were made in R software v4.2.0 (23), using the *ggplot2* package (v3.4.4; Wickham 2016) and *maps* package (v3.4.1.1; Becker and Wilks 2022). Hierarchical clustering of the feeding patterns was done with the *hclust* function from the stats package (v3.6.2) in R utilizing the default settings. To assess the difference in feeding patterns between realms Pearson’s chi-square test was used, no multiple testing correction was applied to the P values. This analysis was performed with the stats package (v3.6.2) in R.

#### Phylogenetic tree

Multiple sequence alignments were made of the cytochrome *c* oxidase subunit I (COI) gene. This gene is commonly used for mosquito species identification as it is easy to amplify and has enough variation in its sequence to distinguish between different species (26). The sequences were obtained from NCBI (National Centre for Biotechnology Information) and only the *Culex* species for which a *COI* gene was available were included for further analysis. With the available sequences a multiple sequence alignment was made in MEGA11 v11 (27) using the ClustalW algorithm. *Chaoboridae* was included as the outgroup. Gblocks 0.91b was utilized to remove poorly covered regions (28). Accession numbers for all of the included species can be found in the supporting information (S3 table). A phylogeny was then created utilizing maximum likelihood methods and 1000 ultrafast bootstraps in IQ-TREE v. 1.6.10 (29).

#### Tanglegram

For the 56 species that had *COI* genes available on NCBI a tanglegram was created which compared *Culex* phylogeny to a tree based on the feeding patterns of the same mosquito. For this tanglegram the Baker’s gamma index (BGI) (30) and the cophenetic correlation coefficient (CCC) (31) was calculated, which give an estimation of the association of the two dendrograms compared in the tanglegram. This tanglegram and the statistical tests were done with the *dendextend* package in R (31).

## Results

### Spatial distribution of *Culex* blood-feeding publications

In total 90 publications were identified that fit the criteria of this study with blood-meals for 71 different *Culex* mosquitoes described. Data on 26,857 blood-meals were recorded. The publications contained *Culex* blood-feeding information from 30 different countries. The United States had both the highest number of publications and the highest number of species described in these publications (<u>Fig</u> 1).

**Fig 1.**
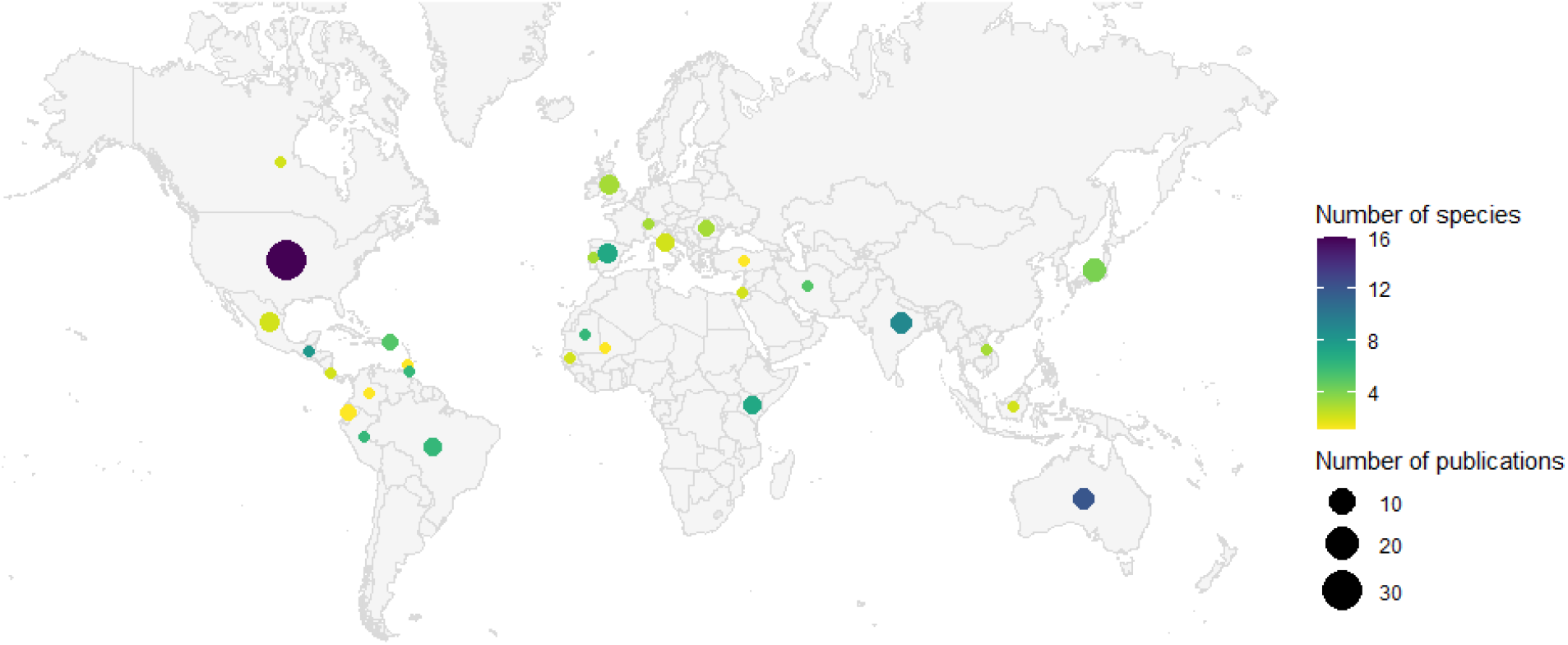
Spatial distribution of *Culex* blood-feeding publications. Circle size represents the number of publications per country. Colour of the circle represents the number of *Culex* species that were analysed per country. The map was created using the *maps* package (v3.4.1.1; License: GPL-2; Becker and Wilks 2022) in *R software* (23).

### Comparison between phylogeny and feeding patterns

A total of 56 *Culex* species were included in the comparison between phylogeny and feeding patterns. Based on the clustering of feeding patterns six broad clusters emerged: majority non-human mammal feeds (*Cx. annulioris, Cx. annulirostris, Cx. caudelli, Cx. coronator, Cx. crinicauda, Cx. declarator, Cx. infula, Cx. poicilipes, Cx. portesi, Cx. salinarius, Cx. spissipes, Cx. taeniopus, Cx. tritaeniorhynchus, Cx. vishnui, Cx. vomerifer*), majority avian feeds (*Cx. australicus, Cx. globocoxitus, Cx. interrogator, Cx. intrincatus, Cx. lactator, Cx. modestus, Cx. mollis, Cx. pip. pallens, Cx. pipiens sp*., *Cx. restuans, Cx. sasai, Cx. stigmatosoma, Cx. territans, ‘Cx. pipiens pooled’*), mix of mammalian and avian feeds (*Cx. erraticus, Cx. erythrothorax, Cx. nigripalus, Cx. orbostiensis, Cx. orientalis, Cx. perexiguus, Cx. pip. molestus, Cx. quinquefasciatus, Cx. restrictor, Cx. sitiens, Cx. tarsalis, Cx. theileri, Cx. torrentium*), majority amphibian feeds (*Cx. amazonensis, Cx. melanoconion sp*., *Cx. microculex sp*.), majority human feeds (*Cx. antennatus, Cx. decens, Cx. dunni, Cx. neavei, Cx. pedroi, Cx. pip. pipiens, Cx. univittatus, Cx. vaxus*), and majority reptile feeds (*Cx. pilosus, Cx. bahamensis, Cx. hortensis*). The phylogenetic clustering and the feeding clustering showed a very low correlation (BKG = 0.27, CCC = 0.17). *Culex vishnui and Culex tritaeniorhynchus* is the only cluster of mosquitoes which cluster in both their phylogeny and feeding patterns (<u>Fig</u> 2).

**Fig 2.**
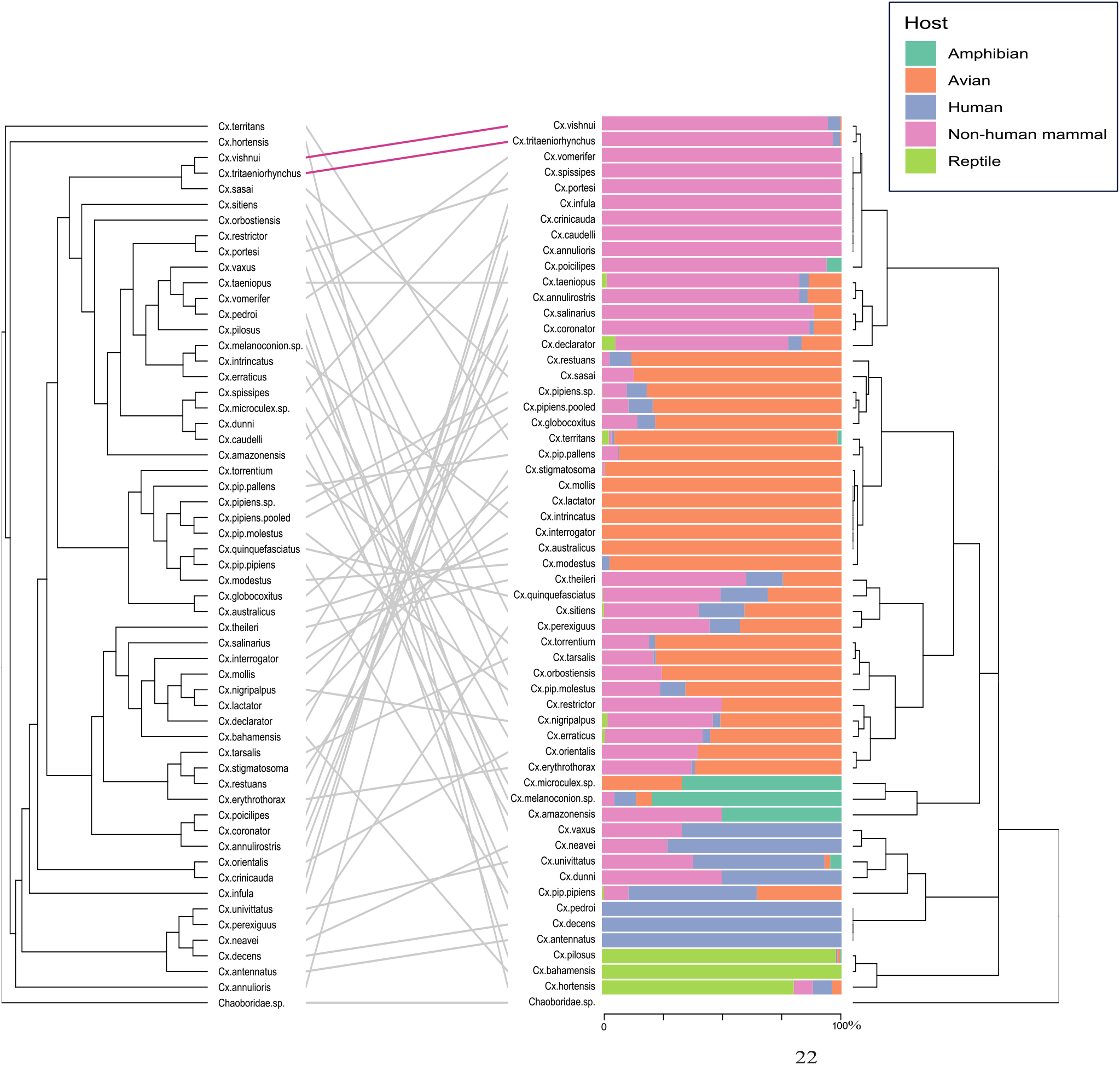
Tanglegram of a phylogenetic tree based on the cytochrome *c* oxidase subunit I (COI) gene of 56 *Culex* (*Cx*.) mosquitoes compared to a hierarchical clustering tree of *Culex* mosquito feeding preferences. The phylogenetic tree (shown on the left) was generated using IQ-TREE with 1000 ultrafast bootstrap alignments. The hierarchical clustering tree (shown on the right) was created in R using the *hclust* function. The height of the trees represents the distance between clusters. Clusters that are the same in both trees are connected with pink lines. Next to the feeding dendrogram the percentage of feeds on each host group is presented per *Culex* species. *Chaoboridae* was chosen as the outgroup. ‘*Culex pipiens’* pooled consist of the following mosquitoes from the *Cx. pipiens* complex: *Culex pipiens, Culex pipiens molestus, Culex pipiens pallens, Culex pipiens pipiens, Culex pipiens/molestus hybrid, Culex pipiens/torrentium*.

**Fig 3.**
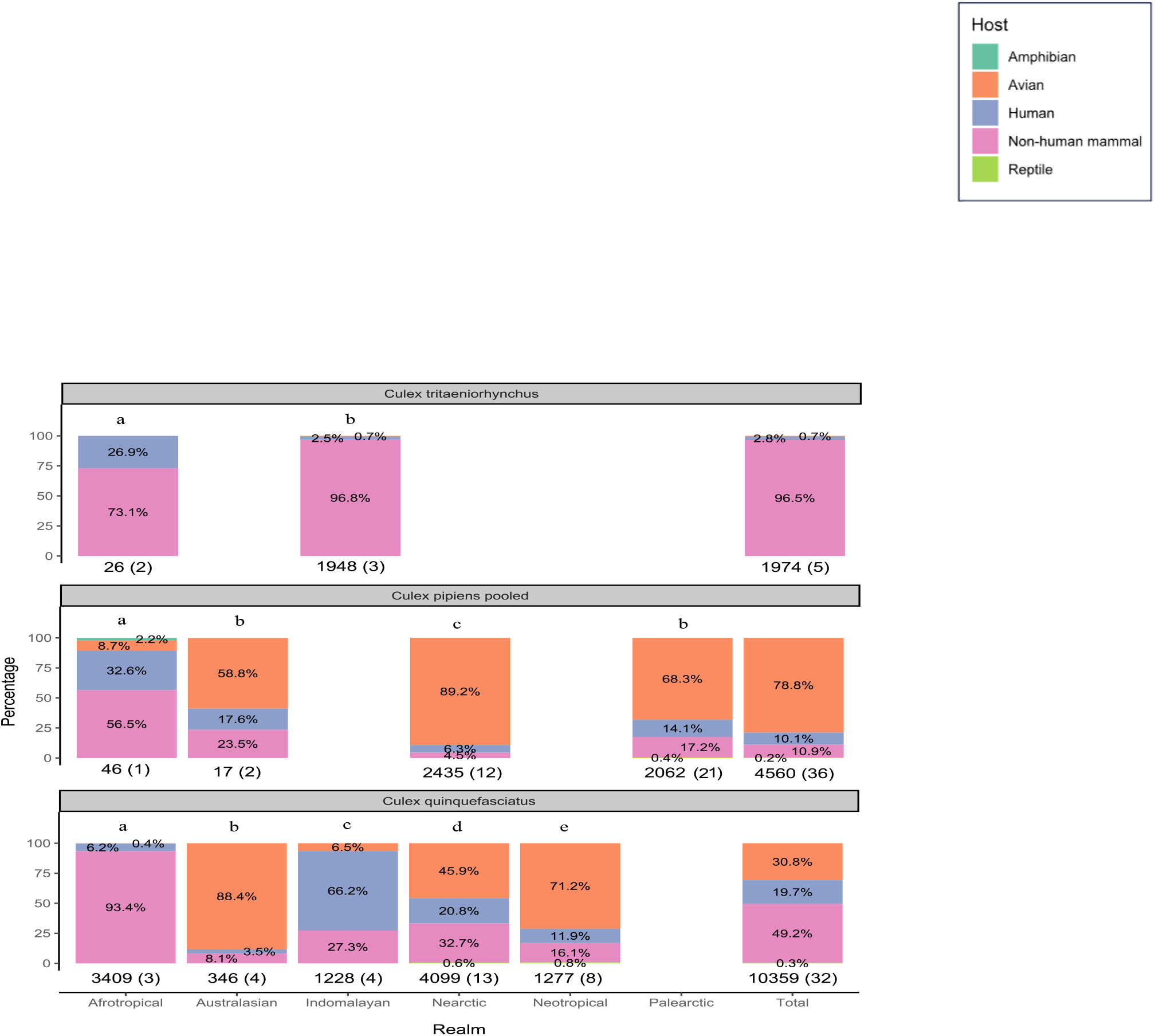
Feeding patterns of *Culex tritaeniorhynchus* (A), *‘Culex pipiens* pooled’ (B), and *Culex quinquefasciatus* (C) in different biogeographical realms. Percentage of blood-meals on amphibian, avian, human, non-human mammal, and reptile hosts are shown for each realm. The total number of blood-meals and total number of publications (blood-meals(publications)) is shown above each bar. ‘*Culex pipiens* pooled’ consist of the following mosquitoes: *Culex pipiens, Culex pipiens molestus, Culex pipiens pallens, Culex pipiens pipiens, Culex pipiens/molestus hybrid, Culex pipiens/torrentium*. Significance groups of Chi-square pairwise comparisons are shown in the figure. If two bars share the same letter they do not differ significantly (p>0.05). If two bars share a different letter they do differ significantly (p<0.05). Subscript numbers separate different pairwise comparisons within the three vector species (i.e. there were no pairwise comparisons between different mosquito species). See supporting information table 4 for full list of pairwise comparisons and P values.

### Feeding patterns across biogeographical realms

*Culex tritaeniorhynchus, ‘Cx. pipiens* pooled’, and *Cx. quinquefasciatus* were chosen for further analysis by realm as they are important vector species which had feeding pattern data available in two or more realms. *Culex tritaeniorhynchus* feeding patterns in the Afrotropical and Indomalayan realm, had a majority of feeds being on non-human mammalian hosts. ‘*Culex pipiens* pooled’ had a majority of avian feeds in the Nearctic, Palearctic and Australasian realms. However, for the Afrotropical realm nearly 90% of *Culex pipiens* pooled feeds was recorded on non-human mammals and humans. All of the realms differed significantly in their feeding patterns for ‘*Culex pipiens* pooled’ except for the Australasian and Palearctic realm (χ2= 2.47, df = 3, p > 0.05). *Culex quinquefasciatus* had a majority of mammal feeds in the Afrotropical realm, human feeds in the Indomalayan realm, and avian feeds in the Australasia, Nearctic, and Neotropical realms. *Culex quinquefasciatus* had significantly different feeding patterns in all included realms. See supporting information table 4 for a full list of pairwise comparisons and P values.

## Discussion

Our analysis of *Culex* feeding patterns revealed that *Culex* mosquitoes have a wide variety of feeding patterns, with some feeding on only mammals, only birds, or only reptiles, but most of them also feeding on multiple host groups. We show that the feeding patterns do not appear to be explained by phylogeny alone. For *Culex pipiens* pooled, *Culex quinquefasciatus, and Culex tritaeniorhynchus* different biogeographical realms showed significantly different feeding patterns. Assessing the feeding patterns of these mosquitoes is essential as they are primary vectors for several important mosquito-borne viruses, including zoonotic viruses (3,4).

*Culex* mosquitoes with a wide host range utilization can potentially act as bridge vectors, e.g. for West Nile virus (WNV), transmitting virus from reservoir birds to other susceptible-but-dead-end hosts such as humans and horses. For example, the competent WNV vectors *Culex restuans, Cx. pipiens sp*., *Cx. theileri, Cx. quinquefasciatus*, and *Cx. modestus* all had feeding patterns that included both human and avian bloodfeeding (32–35). Other studies have described *Cx. restuans* and *Culex pipiens sp*. to be largely ornithophilic (36), which is in line with our results, with 88% of *Cx. restuans* and 81% of *Cx. pipiens sp*. blood-meals being from avian hosts. *Culex quinquefasciatus* is often believed to be a mosquito that can feed on a wide variety of hosts, depending on the abundance of these hosts (37). This is also similar to the results we found as around 31% of all *Cx. quinquefasciatus* feeds was on avian hosts, 20% on humans, 49% on non-human mammals. Very little research has been done on *Cx. modestus* despite this species being a known highly competent vector for WNV in the lab (38). Our results show *Cx. modestus* feeds being 97% on avian hosts. Unpublished observations from the UK seem to support high human biting behaviour (UKHSA, personal communication). The combination of feeding on both birds and humans emphasizes how this species must not be overlooked as a potential WNV vector.

Hierarchical clustering of *Culex* feeding patterns revealed a different grouping than the clusters of a *Culex* phylogeny based on the *COI* gene. No relationship between *Culex* phylogeny and feeding patterns was observed. This is in line with the results from another meta-analysis which summarized the feeding patterns of Australian mosquitoes (39). However, contrasting results were found in another study looking at the feeding patterns of 256 mosquitoes from multiple genera. Their results indicate that evolutionary relationships and major continental drift events are associated with determining feeding patterns (40). Our results only focus on one genus and recent feeding patterns, and do not show an association between *Culex* evolutionary relationships and host feeding patterns. By focussing on one genus, there is no influence of distantly related genera creating clusters (40). Our genus-specific analysis highlights the differences in feeding patterns at a higher resolution, showing that at genus-level analysis, species clusters do not fully explain feeding patterns.

For three geographically widespread and important vector species, we analysed feeding patterns across different biogeographical realms. *Cx. quinquefasciatus* and ‘*Cx. pipiens* pooled’ we observed different host utilizations in different realms. Conversely, for *Cx. tritaeniorhynchus* we saw similar feeding patterns across the two realms it populates. This indicates that the presence of the same species can represent a different risk for different areas. For example, whilst *Cx. quinquefasciatus* might pose a limited threat to humans in the Australasian realm, it could be an important bridge vector in the Afrotropical and Nearctic realm. The reason behind these differences in feeding patterns is unknown, but multiple factors such as host availability, land use, and climate could influence this effect.

This difference in feeding patterns across realms could be due to a different host availability. Since we did not have data on the availability of host species in each specific study area, we cannot determine whether a high percentage of feeds on a specific host species was due to an actual preference of the mosquito for that host or if it was due to that host species being the most available host compared to other species. A more sophisticated way to measure this, if the data were available, would be by utilizing a forage ratio analysis where the observed frequency of blood-meals on a specific host is divided by the relative abundance of that host in the study area (41). Including an estimate of host availability in future field studies would allow for this analysis of feeding patterns. This could be done using regional host density data where it exists.

Land use could be an important factor in determining host utilization of these mosquitoes. Geographically distinct populations of mosquitoes can have different host preferences (42). Urbanization affects feedings patterns as well, with *Cx. pipiens* and *Cx. quinquefasciatus* having increased levels of human feeding in urban areas (12). Additionally, whether the collection occurs inside or outside can have an effect, with inside collections having a significantly higher percentage of human blood feeding compared to collection that just occurred outside (43). The difference in feeding patterns could also be influenced by climate. The intensity of the dry season significantly impacts *Aedes aegypti* preference of human over animal odour (11). Only a few studies make data on the collection location and the surrounding area available which results in a limited insight into the effect of land use in feeding patterns. Future studies should aim to include a description of land use types in a 1 km radius around the collection point of the bloodfed mosquito. If the data are not available studies should provide additional information and/or supplementary pictures of the location of the traps.

Shifts in mosquito feeding behaviour could impact the virus transmission risk to humans. In the US a shift in feeding patterns was observed in *Culex pipiens* after their main host American robins migrated, which resulted in a switch to humans as their primary host. This increase in *Culex pipiens* feeding on humans coincided with an increase in people being infected with West Nile virus (WNV). A similar pattern was observed for *Cx. tarsalis* where an increased number of human WNV infections occurred if the preferred avian host migrated (44). Thus, as land use change and climate change could drive mosquitoes to feed more on human hosts (11,12), this could increase the potential of virus transmission.

When looking at the differences in feeding patterns between realms there was a large variation in the number of blood-meals recorded in each realm. Additionally, the number of publications available for each species varied greatly. As we did not correct for sampling effort the results might be biased towards feeding patterns that are common in more well sampled areas. Additionally, for some species complexes that are challenging and inconsistent to identify, such as the *Culex pipiens* complex, data on individual sub-species/biotype are lacking in many included studies. Consequently, our analyses may not accurately reflect the feeding patterns of lower-order taxa e.g. sub-species for this complex. For most of the studies the local host availability, season, land use, exact time and location of the collected mosquito was not recorded. These details would have given a more comprehensive understanding of the mosquito host feeding patterns. *Culex pipiens* pooled showed variation in feeding patterns between realms. The different subtypes contained in this pool have been shown to have different feeding preferences. *Culex pipiens pipiens* and *Culex pipiens molestus* are an example of this, with biotype *pipiens* having a high percentage of avian blood-meals whilst biotype *molestus* had a higher percentage of human blood-meals (45). Thus, the differences in feeding patterns between the realms for *Culex pipiens* pooled could have been due to the different *Culex pipiens* subtypes contained in the pool being mostly present in different locations.

Analysing the feeding patterns of *Culex* mosquitoes on a global scale showed a large variety in feeding patterns between the *Culex* species and a limited genetic basis for these differences. Additionally, variations in feeding patterns in the different realms for the same species were found, highlighting the importance of local surveillance. The reason behind these different patterns is unknown but indicates extrinsic factors such as host availability, land use and climate playing a large role. When determining how host feeding patterns influence human disease risk a wide variety of external factors should therefore be included in the consideration. Finally, as climate and land use change may alter the feeding patterns on avian and human hosts, there may be an increased potential of mosquito-borne viral epidemics and cross species transmission. There are not enough data available to predict this shift and its possible effects at present, again highlighting the importance of local and periodic surveillance in high-risk areas.

## Supporting information

**S1 Table. Full dataset of all included studies**

**S2 Code. Code utilized for the meta-analysis of *Culex* blood-feeding studies**

**S3 Table. Accession numbers of mosquitoes utilized to create phylogenetic tree**

**S4 Table. Full statistical analysis of feeding across realms**

## References

1. Farajollahi A, Fonseca DM, Kramer LD, Marm Kilpatrick A. “Bird biting” mosquitoes and human disease: A review of the role of Culex pipiens complex mosquitoes in epidemiology. Infect Genet Evol [Internet]. Elsevier B.V.; 2011;11(7):1577–85. Available from: 10.1016/j.meegid.2011.08.013

2. Harbach RE. Classification within the cosmopolitan genus Culex (Diptera: Culicidae): The foundation for molecular systematics and phylogenetic research. Acta Trop [Internet]. Elsevier B.V.; 2011;120(1–2):1–14. Available from: 10.1016/j.actatropica.2011.06.005

3. Bernard KA, Kramer LD. West Nile virus activity-United States, 2001. Viral Immunol. 2001;14(4):319–38.

4. Van Den Hurk AF, Ritchie SA, Mackenzie JS. Ecology and geographical expansion of japanese encephalitis virus. Annu Rev Entomol. 2009;54:17–35.

5. Vloet RPM, Vogels CBF, Koenraadt CJM, Pijlman GP, Eiden M, Gonzales JL, et al. Transmission of Rift Valley fever virus from European-breed lambs to Culex pipiens mosquitoes. PLoS Negl Trop Dis. 2017;11(12):1–17.

6. Calzolari M, Bonilauri P, Bellini R, Albieri A, Defilippo F, Maioli G, et al. Evidence of simultaneous circulation of west Nile and Usutu viruses in mosquitoes sampled in Emilia-Romagna region (Italy) in 2009. PLoS One. 2010;5(12):1–10.

7. Dunphy BM, Kovach KB, Gehrke EJ, Field EN, Rowley WA, Bartholomay LC, et al. Long-term surveillance defines spatial and temporal patterns implicating Culex tarsalis as the primary vector of West Nile virus. Sci Rep [Internet]. Springer US; 2019;9(1):1–10. Available from: 10.1038/s41598-019-43246-y

8. Kim H, Cha GW, Jeong YE, Lee WG, Chang KS, Roh JY, et al. Detection of Japanese encephalitis virus genotype V in Culex orientalis and Culex pipiens (Diptera: Culicidae) in Korea. PLoS One. 2015;10(2):1–13.

9. Mukabana WR, Takken W, Knols BGJ. Analysis of arthropod bloodmeals using molecular genetic markers. Trends Parasitol. 2002;18(11):505–9.

10. Kilpatrick AM, Kramer LD, Campbell SR, Alleyne EO, Dobson AP, Daszak P. West Nile virus risk assessment and the bridge vector paradigm. Emerg Infect Dis. 2005;11(3):425–9.

11. Rose NH, Sylla M, Badolo A, Lutomiah J, Ayala D, Aribodor OB, et al. Climate and Urbanization Drive Mosquito Preference for Humans. Curr Biol [Internet]. Elsevier Ltd.; 2020;30(18):3570–3579.e6. Available from: 10.1016/j.cub.2020.06.092

12. Santiago-alarcon D. Ecology of diseases transmitted by mosquitoes to wildlife. Ecology of diseases transmitted by mosquitoes to wildlife. 2022. 161–177 p.

13. Ulloa A, Arredondo-Jimenez J, Rodriguez MH, Fernandez-salas I, Gonzalez-ceron L. Innate host selection in Anopheles vestitipennis from southern Mexico. J Am Mosq Control Assoc. 2004;20(4):337–41.

14. Huang S, Hamer GL, Molaei G, Walker ED, Goldberg TL, Kitron UD, et al. Genetic variation associated with mammalian feeding in Culex pipiens from a West Nile virus epidemic region in Chicago, Illinois. Vector-Borne Zoonotic Dis. 2009;9(6):637–42.

15. Thiemann TC, Wheeler SS, Barker CM, Reisen WK. Mosquito host selection varies seasonally with host availability and mosquito density. PLoS Negl Trop Dis. 2011;5(12).

16. Takken W, Verhulst NO. Host preferences of blood-feeding mosquitoes. Annu Rev Entomol. 2013;58(September):433–53.

17. MacKay AJ, Kramer WL, Meece JK, Brumfield RT, Foil LD. Host feeding patterns of culex mosquitoes (Diptera: Culicidae) in east baton rouge Parish, Louisiana. J Med Entomol. 2010;47(2):238–48.

18. Kreft H, Jetz W. A framework for delineating biogeographical regions based on species distributions. J Biogeogr. 2010;37(11):2029–53.

19. Calzolari M. Mosquito-borne diseases in Europe: an emerging public health threat. Reports Parasitol [Internet]. 2016;1. Available from: 10.2147/RIP.S56780

20. Semenza JC, Paz S. Climate change and infectious disease in Europe: Impact, projection and adaptation. Lancet Reg Heal - Eur [Internet]. Elsevier Ltd; 2021;9:100230. Available from: 10.1016/j.lanepe.2021.100230

21. Soto A, Delang L. Culex modestus: the overlooked mosquito vector. Parasites and Vectors [Internet]. BioMed Central; 2023;16(1):1–14. Available from: 10.1186/s13071-023-05997-6

22. Udvardy MDF. A Classification of the Biogeographical Provinces of the World. Union Conserv Nat Nat Resour. 1975;

23. R Core Team. R: A language and environment for statistical computing [Internet]. Vienna, Austria: Foundation for Statistical Computing; 2022. Available from: https://www.r-project.org/.

24. Wickham H. ggplot2 - Elegant Graphics for Data Analysis (2nd Edition). J Stat Softw. 2016;

25. Becker RA, Wilks AR. Draw Geographical Maps. 2022;

26. Chan A, Chiang LP, Hapuarachchi HC, Tan CH, Pang SC, Lee R, et al. DNA barcoding: Complementing morphological identification of mosquito species in Singapore. Parasites and Vectors. 2014;7(1):1–12.

27. Tamura K, Stecher G, Sudhir K. MEGA11: Molecular Evolutionary Genetics Analysis version 11. Mol Biol Evol. 2021;38:3022–7.

28. Castresana J. Selection of conserved blocks from multiple alignments for their use in phylogenetic analysis. Mol Biol Evol. 2000;17(4):540–52.

29. Nguyen LT, Schmidt HA, Von Haeseler A, Minh BQ. IQ-TREE: A fast and effective stochastic algorithm for estimating maximum-likelihood phylogenies. Mol Biol Evol. 2015;32(1):268–74.

30. Baker FB. Stability of Two Hierarchical Grouping Techniques Case I: Sensitivity to Data Errors. J Am Stat Assoc. 1972;69(346):440–5.

31. Galili T. dendextend: An R package for visualizing, adjusting and comparing trees of hierarchical clustering. Bioinformatics. 2015;31(22):3718–20.

32. Ebel GD, Rochlin I, Longacker J, Kramer LD. Culex restuans (Diptera: Culicidae) relative abundance and vector competence for West Nile virus. J Med Entomol. 2005;42(5):838–43.

33. Shahhosseini N, Moosa-kazemi SH, Sedaghat MM, Nowotny N, Kayedi MH. Autochthonous Transmission of West Nile Virus by a New Vector in Iran, Vector-Host Interaction Modeling and Virulence Gene Determinants. Viruses. 2020;12(12).

34. Eastwood G, Kramer LD, Goodman SJ, Cunningham AA. West Nile virus vector competency of Culex quinquefasciatus mosquitoes in the Galápagos Islands. Am J Trop Med Hyg. 2011;85(3):426–33.

35. Hubálek Z, Halouzka J. West Nile fever - A reemerging mosquito-borne viral disease in Europe. Emerg Infect Dis. 1999;5(5):643–50.

36. Molaei G, Andreadis TG, Armstrong PM, Anderson JF, Vossbrinck CR. Host feeding patterns of Culex mosquitoes and west nile virus transmission, northeastern United States. Emerg Infect Dis. 2006;12(3):468–74.

37. Molaei G, Andreadis TG, Armstrong PM, Bueno R, Dennett JA, Real S V., et al. Host feeding pattern of Culex quinquefasciatus (Diptera: Culicidae) and its role in transmission of West Nile virus in Harris County, Texas. Am J Trop Med Hyg. 2007;77(1):73–81.

38. Balenghien T, Vazeille M, Grandadam M, Schaffner F, Zeller H, Reiter P, et al. Vector competence of some French Culex and Aedes mosquitoes for West Nile Virus. Vector-Borne Zoonotic Dis. 2008;8(5):589–95.

39. Stephenson EB, Murphy AK, Jansen CC, Peel AJ, McCallum H. Interpreting mosquito feeding patterns in Australia through an ecological lens: An analysis of blood meal studies. Parasites and Vectors [Internet]. BioMed Central; 2019;12(1):1–11. Available from: 10.1186/s13071-019-3405-z

40. Soghigian J, Sither C, Justi SA, Morinaga G, Cassel BK, Vitek CJ, et al. Phylogenomics reveals the history of host use in mosquitoes. Nat Commun. Springer US; 2023;14(1).

41. Komar N, Panella NA, Golnar AJ, Hamer GL. Forage Ratio Analysis of the Southern House Mosquito in College Station, Texas. Vector-Borne Zoonotic Dis. 2018;18(9):485–90.

42. Williams CR, Kokkinn MJ, Smith BP. Intraspecific variation in odor-mediated host preference of the mosquito Culex annulirostris. J Chem Ecol. 2003;29(8):1889–903.

43. Orsborne J, Furuya-Kanamori L, Jeffries CL, Kristan M, Mohammed AR, Afrane YA, et al. Using the human blood index to investigate host biting plasticity: A systematic review and meta-regression of the three major African malaria vectors. Malar J [Internet]. BioMed Central; 2018;17(1):1–8. Available from: 10.1186/s12936-018-2632-7

44. Kilpatrick AM, Kramer LD, Jones MJ, Marra PP, Daszak P. West Nile virus epidemics in North America are driven by shifts in mosquito feeding behavior. PLoS Biol. 2006;4(4):606–10.

45. Osório HC, Zé-Zé L, Amaro F, Nunes A, Alves MJ. Sympatric occurrence of Culex pipiens (Diptera, Culicidae) biotypes pipiens, molestus and their hybrids in Portugal, Western Europe: Feeding patterns and habitat determinants. Med Vet Entomol. 2014;28(1):103–9.

